# Using Eye Gaze to Train an Adaptive Myoelectric Interface

**DOI:** 10.1101/2024.04.08.588608

**Authors:** Amber H.Y. Chou, Maneeshika Madduri, Si Jia Li, Jason Isa, Andrew Christensen, Finley (Liya) Hutchison, Samuel A. Burden, Amy L. Orsborn

## Abstract

Myoelectric interfaces hold promise in consumer and health applications, but they are currently limited by variable performance across users and poor generalizability across tasks. To address these limitations, we consider interfaces that continually adapt during operation. Although current adaptive interfaces can reduce inter-subject variability, they still generalize poorly between tasks because they make use of task-specific data during training. To address this limitation, we propose a new paradigm to adapt myoelectric interfaces using natural eye gaze as training data. We recruited 11 subjects to test our proposed method on a 2D computer cursor control task using high-density surface EMG signals measured from forearm muscles. We find comparable task performance between our gaze-trained paradigm and the current task-dependent method. This result demonstrates the feasibility of using eye gaze to replace task-specific training data in adaptive myoelectric interfaces, holding promise for generalization across diverse computer tasks.

**CCS Concepts:** **• Human-centered computing → Interaction devices; Empirical studies in HCI.**

## 1 INTRODUCTION

Interfaces that use physiological inputs – biosignal-based interfaces [41] – are increasingly popular in consumer and health applications due to the rich human-computer interactions they can facilitate across diverse populations [6, 7, 65, 70]. One class of interfaces converts surface electromyographic (sEMG) activity, which are electrical signals generated by muscle contractions and recorded at the skin, into control commands for external devices such as a computer cursor or a bionic limb [18, 19]. These *myoelectric interfaces* have strong accessibility potential as they are non-invasive and can provide high-dimensional recordings with high control bandwidth from a variety of muscles [4, 18, 70]. However, there are two fundamental challenges in the design of myoelectric interfaces: sEMG signals show large variability between subjects and within a subject over time [4, 67], and the mapping from myographic activity to task variables may be unintuitive.

To address the challenges of signal variability and interface complexity, it is natural to apply machine learning. For instance, offline methods can be used to synthesize an interface to optimize performance with respect to the distribution of subjects represented in a set of labeled training data [39, 50, 60, 75, 76]. However, the performance observed online with this approach consistently falls short of that obtained during offline training [9, 15, 27, 38]. This finding is unsurprising, since users are not stationary data sources – they adapt to the interfaces they use and tasks they are assigned. Adapting the interface online can potentially close this train/test performance gap [23, 27, 41, 69], but presents a new challenge: how to obtain labeled training data at the same time that the interface is being used? Current adaptive algorithms address this challenge by making use of information about the tasks prescribed to users [25, 42, 51], but requiring this information limits generalizability of the interface.

Our vision is to create personalized interfaces that continuously adapt while users perform a variety of tasks, including less structured and self-paced tasks like drawing or internet browsing. Toward this aim, we investigated the use of *eye gaze* as a new task-agnostic source of training data labels for adaptive myoelectric interface algorithms (Fig. 1a). Conventional gaze-controlled interfaces generally require users to focus their gaze to produce input, which limits natural eye movements during everyday tasks and can lead to fatigue [34]. Natural eye behavior, in contrast, offers insights into user’s movement goals [35, 46], which has the potential to augment human-computer systems with indirect signals [3]. Therefore, we hypothesized that eye movements can replace task-based data in training adaptive myoelectric interfaces by assuming that natural gaze serves as a proxy for user intent and task goals. This gaze-trained paradigm removes the constraint of task-based supervision in adaptive algorithms, potentially enabling generalization across diverse tasks. To our knowledge, this work has the first adaptive myoelectric interface trained online using eye gaze data.

**Fig. 1.**
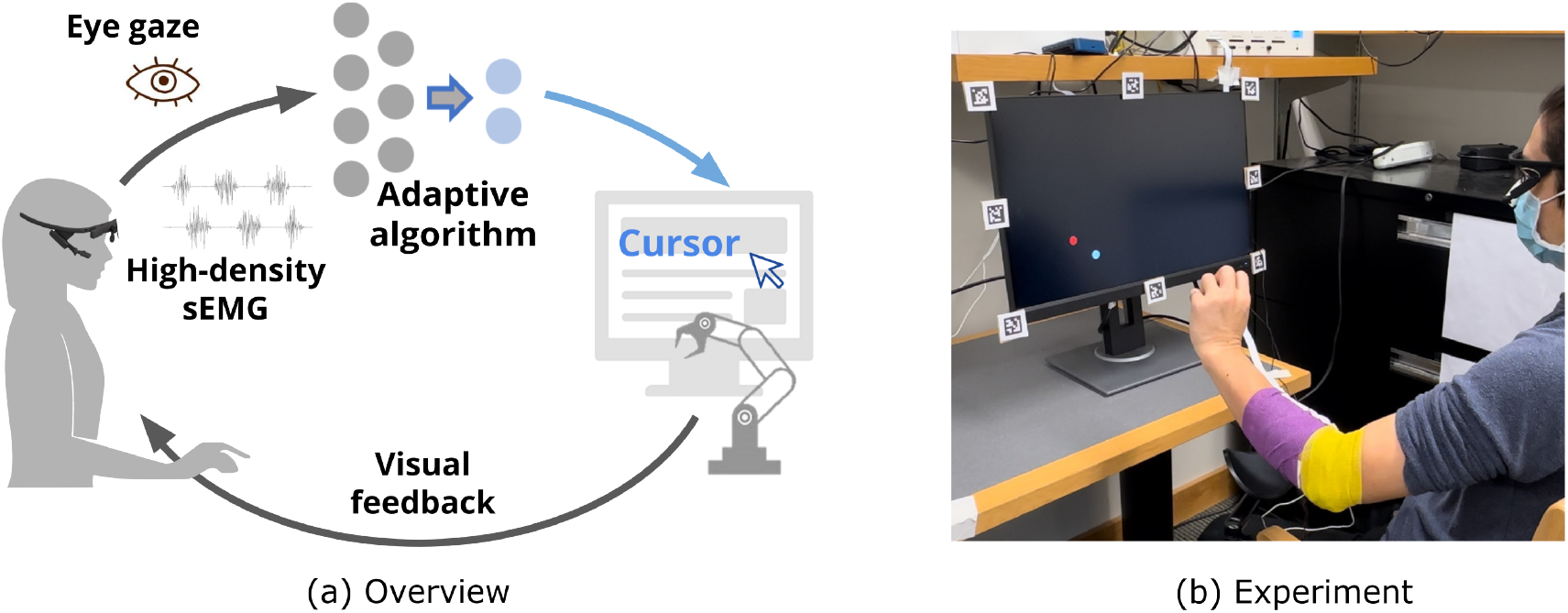
(a) Overview of a closed-loop myoelectric interface that uses eye gaze for training an adaptive mapping to control an external device, such as a computer cursor or a robotic arm. (b) The user controls a cursor (white) to track a continuous target (red) on a computer screen using high-density surface electromyography (sEMG) on the forearm. Gaze data is collected with an eye-tracking headset and is used to train the adaptive algorithm.

To demonstrate the feasibility of the new gaze-trained paradigm, we recruited 11 naive subjects to control a computer cursor using a 64-channel sEMG electrode on their forearm to follow a 2-dimensional (2D) moving target (Fig. 1b). Previous literature defined this cursor-tracking as a *continuous task* [15, 70], which has the potential to be translated to other continuous computer or human-device tasks, such as handwriting [32, 39], prostheses control [21, 75], or wheelchair navigation [49]. Our results demonstrate the practical use of our gaze-trained paradigm, which worked as effectively as the previous task-trained method when natural eye movements were present.

The specific contributions of our work are twofold:

1. We propose a new training paradigm to adapt myoelectric interfaces using eye gaze as training data and highlight its task-agnostic potential for diverse computer tasks.
2. We show that our gaze-trained paradigm has the same effective performance as the prior task-dependent method both with and without guided instruction to users regarding where to fixate their gaze.

This study is an initial step in developing an out-of-the-box myoelectric device that seamlessly self-calibrates to diverse users and tasks. The findings of this study can benefit the development of wearables in HCI applications, particularly in everyday tasks like continuous cursor control in computers or extended reality environments.

## 2 RELATED WORK

### 2.1 Current Myoelectric and Adaptive Algorithms

Myoelectric interfaces translate high-dimensional muscle activity to low-dimensional control of an external device. The process involves recording and filtering raw sEMG activity, followed by the selection of relevant features [16]. These selected features are then mapped to low-dimensional control signals, commonly achieved through linear dimensionality reduction (e.g, principal component analysis or non-negative matrix factorization [13, 50]), regression [27, 41], or classification [48, 60]. These mappings in myoelectric interfaces are typically programmed offline, without the user in the loop [13]. For instance, prior to operating an interface, users’ muscle activity is measured to capture specific movements or contractions, and this recorded activity is used to program a fixed mapping. Although this scheme has promising offline accuracy, the dynamic nature of human users in the loop and the large signal variability in sEMG [67] often results in a mismatch between offline algorithm performance and real-time interface usage [15, 27, 66]. Another limitation of myoelectric interfaces in current studies is the pre-assigning gestures for specific tasks. Many gesture recognition-based algorithms require users to learn a specific set of gestures and then map those pre-set gestures to device commands [8, 28, 48, 60]. However, users have highly varied and personalized gestures that they prefer to use [69]. Thus, pre-assigning gestures limit the flexibility and accessibility of interfaces.

Many high-dimensional inputs in brain-computer interfaces (BCI), such as intracortical neural activities, have the same limitations as above when operating in a closed loop. To address these limitations, prior BCI studies have demonstrated the effectiveness of adapting the mapping online, that is, adapting while the user is operating the interface. Orsborn et al. [52] used a gradient-based supervised learning algorithm to update the mapping based on real-time neural activity. Madduri et al. [42] expanded the method and tested the algorithm as naive users learned a myoelectric interface. These studies demonstrated that online adaptation provides rapid calibration, improves interface performance, accounts for signal nonstationarities, and individualizes the interface to diverse users [14, 26, 41]. Moreover, adaptive interfaces reduce the reliance on guided instructions, allowing users to customize their gestures and strategies at their own pace. Users can also re-train interfaces on the fly, enhancing long-term usability.

Adaptive interfaces are not new in the HCI community, as they have been studied in many machine learning use cases [15, 17]. However, adoption of adaptive algorithms is still limited in myoelectric applications. This work builds off of an adaptive algorithm explored in prior BCI literature [42, 52] to design an adaptive myoelectric interface that continuously calibrates during operations. This interface is specifically tailored for myoelectric use in HCI applications. For instance, using a wearable device to perform everyday computer tasks like cursor navigation. The novelty of this work is the gaze-trained paradigm that trains the adaptive myoelectric interface using natural gaze. This method allows users to decide their own strategies for myoelectric control without the need for prescribed tasks and gestures, or guided eye movements.

### 2.2 Myoelectric Interfaces in HCI

Noninvasive myoelectric sensing has been explored in human-device interaction studies, including teleoperation [20], user understanding [43], VR/AR training [55], and upper-limb [18] and lower-limb prostheses [21], and rehabilitation [33, 47, 50, 56]. From an HCI perspective, these studies demonstrated an accessible alternative to traditional manual devices like mice and keyboards for users with upper-limb motor impairments [70]. Additionally, myoelectric interfaces provide always-available inputs that enable everyday gesture detection to replace touch-based mobile interfaces [31, 39, 61]. This is particularly desirable when a user’s hands are occupied or when a touch screen is not available.

Particularly, prior HCI studies have shown the promise of using upper body (e.g., forearm) muscle signals as an input modality to a computer device. Saponas et al. [60, 61] and Huang et al. [31] used sEMG armbands to classify hand gestures such as thumb tapping and finger lifting for mobile or keyboard interactions. Others have also studied sEMG in gesture recognition for smartwatch [71] and smart garment [5] interaction. In addition to discrete hand gestures, many studies have demonstrated the use of sEMG for continuous cursor navigation through hand movements or muscle contractions [37, 40, 57, 70]. CTRL-labs at Reality Labs [39] has recently reported an sEMG decoding model trained with thousands of participants’ data for multiple computer tasks like continuous navigation, discrete gestures, and handwriting. They have found that their deep-learning model can be accurately generalized across populations and sessions.

While the interfaces in these studies worked effectively for their specific purposes, they were trained using offline data, which hindered their adaptability across users and tasks when a dynamic user is present in the loop. While CTRL-labs’ findings were exceptional, it is challenging to replicate their model built with such large subject sizes. This work also uses sEMG as an input modality for computer cursor control; however, the main difference between this work and the prior work is the continuous adaptation of the myoelectric interface while users operate a task. In addition, our adaptive myoelectric algorithm does not require a large dataset collected offline; instead, the mapping is trained online with data from a single participant in a trial. This can potentially avoid issues such as replicability or biosignal data privacy.

### 2.3 Eye Tracking in HCI

Eye-tracking technology has found widespread applications in virtual reality headsets to enrich user understanding and create more immersive experiences. In addition, eye-tracking serves as a readily measurable input modality to control devices without hands, and is faster in cursor control compared to conventional mice [34, 72]. Prior studies have shown that eye inputs can augment interface functionalities [34, 74] and enhance interface accessibility [29, 73].

Several studies in HCI have shown the advantages of integrating interfaces with eye-tracking and upper-body movements. For instance, the combination of gaze with gesture inputs enables rapid, touch-free computer interactions [10] and can be used to enhance user intention estimation [62]. Other studies developed multimodal interfaces that used gaze for cursor navigation and sEMG (with facial muscles, forehead, forearm, etc.) for discrete operations such as selection, switching, or clicking, demonstrating performance surpassing that of traditional mice [44, 53]. While combining multiple sources of biosignals enhances interactions, the potential benefits of multiple biosignals in adaptive interfaces have not yet been fully explored.

In this work, we extend previous research on myoelectric interfaces by incorporating eye gaze data. What sets our approach apart from other gaze-controlled devices is that we did not directly use intentional gaze as a control input. Rather, we used natural gaze as training data for our adaptive algorithm. Several studies have also integrated people’s natural gaze in human-robot interaction and have proved its advantages in anticipating user selections or intentions, thus improving system efficiency and task accuracy [1, 3, 11, 30, 63]. Aronson et al. [3] used natural gaze to predict users’ task goals and then used the predictions to improve a learning algorithm for manual (joystick)-based robot manipulation. Similarly, this study also leverages gaze to approximate the user’s desired actions. The main distinction lies in the interface adaptation using high-dimensional biosignal inputs, contrasting with the fixed mapping of low-dimensional manual devices used in previous studies.

## 3 EYE-GAZE FACILITATED ADAPTIVE ALGORITHM

Online adaptation was used to update the matrix *D* that mapped high-dimensional sEMG signals *s* to the velocity *v* of a 2D cursor:

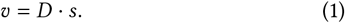

The matrix *D* was adapted using a gradient-based learning algorithm designed to minimize the task error with a regularization term [42]. In prior work that used task information for training the algorithm, it was assumed that the user would intend to move the cursor toward the task target [25, 51]; thus, the intended goal was the target position *r*. The task error was therefore defined as the norm difference between the intended velocity – the instantaneous velocity of a vector between intended goal *r* and cursor *y* at a given time – and the actual velocity of the cursor:

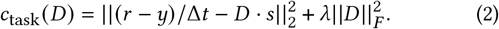

where Δ*t* is the time interval, ||*D* ||_*F*_ denotes the Frobenius norm of the matrix *D* ∈ ℝ^2*xN*^ where *N* is the number of EMG input features, and *λ* is the weight of the regularization.

The innovation of this work is to infer the user’s intended goal from their gaze and use gaze for training the interface. We replaced the intended goal in the cost function to be gaze position *g* in the plane defined by the computer display:

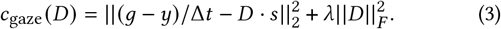

Thus, the adaptive algorithm was not explicitly given the target information in the gaze-trained interface.

Following the method in Madduri et al. [42], the matrix *D* was updated iteratively to minimize the cost (*c*_task_ or *c*_gaze_) using 20-second batches of data:

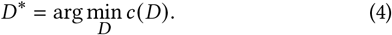

The 20-second batch length was selected to train data with intended velocities spanning various directions on the computer screen.

## 4 METHODS

### 4.1 Participants

We recruited 11 participants without disabilities (5 males, 5 females, 1 non-binary; 2 left-handed, 9 right-handed; aged 19 to 32). They were included in the study if they had normal or corrected-to-normal eyesight and no history of upper-limb disability. Three participants had used an EMG device before, and two participants regularly worked with EMG devices. Participants gave their written consent prior to the study. All procedures were approved by the University of Washington’s Institutional Review Board (IRB #STUDY00014060).

### 4.2 Data Acquisition and Processing

User myoelectric activity was recorded from a high-density sEMG 64-channel electrode (5×13 rectangular electrode layout, 4mm inter-electrode spacing from Quattrocento, Bioelectronica, Italy) that was placed on the user’s dominant forearm, targeting the Extensor Carpi Radialis. The electrode array was wrapped with self-adherent tape. EMG signals were recorded using OT Bio-Light Software (Bioelettronica, Italy) at 2048 Hz on Differential Mode with a built-in high-pass filter at 10 Hz, and a low-pass filter at 150 Hz. Then, raw EMG signals were digitally rectified and low-pass filtered by averaging the delinearized signals with 100 ms time bins [70] to obtain an EMG envelope that was input to the interface algorithm.

The eye data were collected using the PupilLabs Core headset (PupilLabs, Germany) [36]. The gaze data streamed at 250 Hz using the PupilCapture Software (PupilLabs, Germany), combining the streaming data from the two infrared cameras in real time. Eight AprilTags markers were placed around a 24-inch computer monitor to define an area of interest in the PupilCapture Software (Fig. 1b). Calibration in PupilCapture was conducted by a process where users had to gaze at five static targets on the screen. The monitor was 27 inches in front of the participant. The heights of the monitor, table, and chair were adjustable to maximize participants’ comfort level. In the condition where gaze measurements were used online to train the adaptive algorithm, the gaze data in every 20-second batch was filtered to remove low-confidence estimations and de-biased for any calibration errors (see Appendix A).

### 4.3 User Task

The experiment was described to the participants as a continuous trajectory-tracking task. Participants were instructed to control a cursor to stay as close to a 2D moving target as possible at all times. We constructed target references as sums of sinusoidal signals with random phases to create a pseudorandom target trajectory. The sinusoidal signals were designed to be at frequencies of prime multiples of the base frequency (0.05 Hz) below 1 Hz [58, 68]. The signals were at 0.1, 0.25, 0.55, 0.85 Hz in the horizontal direction and 0.15, 0.35, 0.65, 0.95 Hz in the vertical direction. We called these frequencies the *stimulated frequencies* in this paper, denoted as a set Ω. The magnitudes of stimuli were the inverse of the frequencies to distribute signal power equally. The target and cursor positions on the computer screen were updated at 60 Hz.

### 4.4 Conditions and Procedure

At the beginning of each experiment, participants conducted a baseline trial using a computer mouse to perform the continuous tracking task. This baseline trial used the conventional mapping from the mouse position to the cursor position, and this mapping was not adapted.

After the baseline trial, participants performed the same tracking task but now controlled via an adaptive myoelectric interface. They conducted myoelectric interface control with two conditions: **(1) Task-trained:** using target position as training data (Eq. 2), and **(2) Gaze-trained:** using gaze position as training data (Eq. 3). Each trial was three minutes and each condition was repeated four times during this experiment. Participants performed four consecutive blocks in total, each with both conditions. Participants did not know the training mechanism of this experiment, nor which training method was performed. The condition orders were randomized in each block to control for order effects. The mapping *D* was initialized randomly (matrix entries sampled uniformly at random in the range [0, 10^*−*2^]) and then updated every 20 seconds during training.

To understand whether guided instruction on gaze would affect the performance of the gaze-trained interface, we conducted two sessions in this study–each session had two blocks–with a five-minute break in between sessions (Fig. 2a). In *Session 1*, no explicit instructions on the user’s gaze were given by the experimenter, thus natural gaze was used as training input; while in *Session 2*, participants were instructed to gaze at the target as closely as they could in all trials (Fig. 2b).

**Fig. 2.**
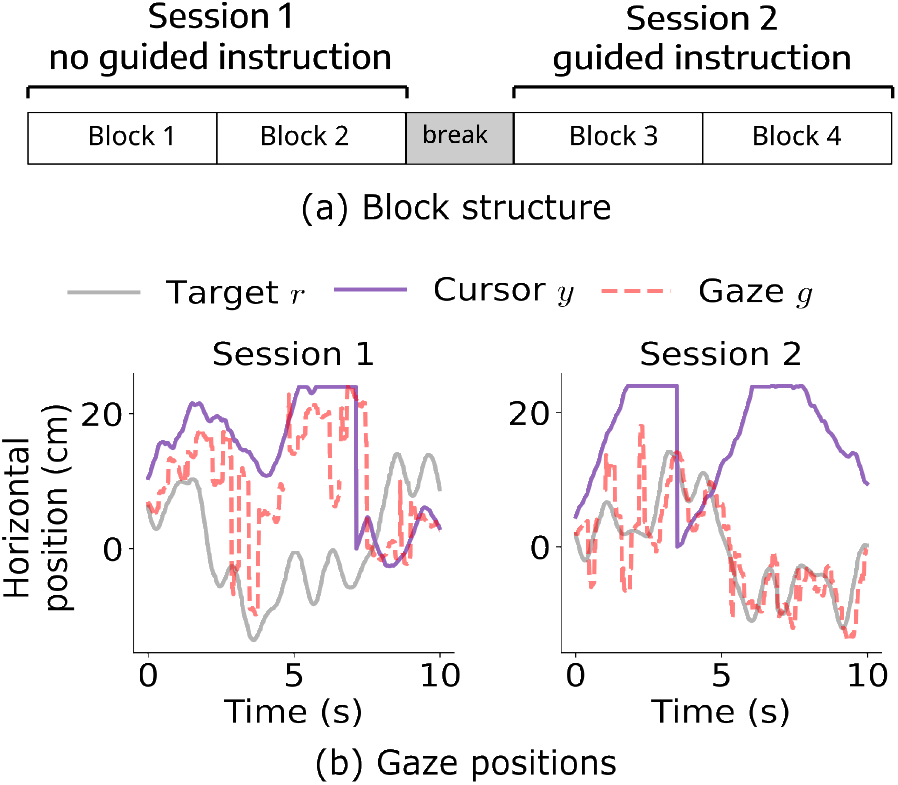
(a) Block structure of the experiment. Participants conducted four blocks and took a mandatory five-minute break between sessions. *Session 2* had guided instruction on where to gaze. (b) Example gaze positions *g* over time in *Session 1* (left) and *Session 2* (right).

At the end of the experiment, participants filled out a brief questionnaire to subjectively quantify their comfort with this system, across three categories: task difficulty, interface accuracy, and effort. We report these subjective results in Appendix B.

### 4.5 Evaluation

To measure the online usability of the adaptive interface, we applied the online evaluation method discussed in prior literature [15] and quantified the continuous task performance as the time-domain tracking ability [70]. Specifically, we quantified task performance over time interval [*t*_0_, *t*_1_] using the mean square error between the 2D target *r* and cursor *y* signals,

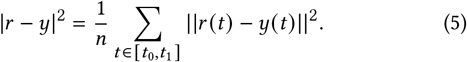

where *n* is the number of time samples in the interval [*t*_0_, *t*_1_]. Lower error corresponds to better task performance.

To assess how performance changed during training, we compared task error from *early* to *late* periods within trials (as illustrated in Fig. 3) across both sessions for each of three conditions: task-trained, gaze-trained, and mouse baseline. Since the mapping was initialized randomly and updated after 20 seconds, we removed the initial 20 seconds of data and compared the subsequent 20 seconds (*early*) with the last 20 seconds (*late*) of adaptation for each condition. The non-adaptive conventional mouse interface was used as a baseline for comparison with performance achieved in the *late* period of trials with our adaptive myoelectric interfaces. We verified normality of the data using the Shapiro-Wilks test (task-trained: *p* = 0.67; gaze-trained: *p* = 0.89; baseline: *p* = 0.80) and applied paired t-tests and repeated measure analysis of variance (ANOVA) test with *α* = 0.05. We used paired t-tests with Bonferroni correction as post-hoc tests. We had the following two hypotheses for interface performance after training:

**Fig. 3.**
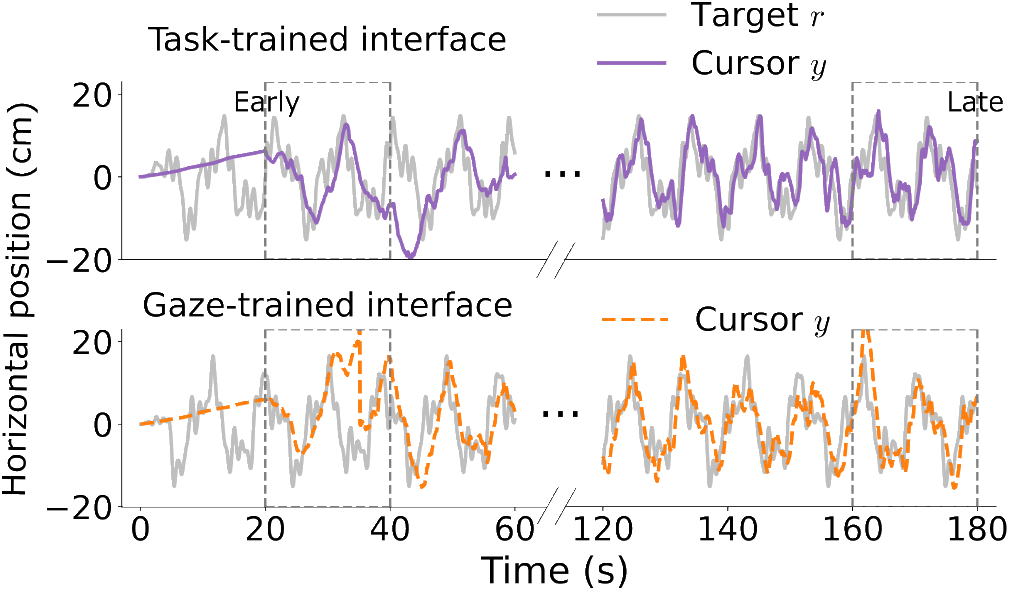
Two example trials of the continuous tracking task. Both task- and gaze-trained interfaces had better tracking performance in the third minute (right) compared to the first minute (left) of a trial. The *early* and *late* 20-second windows are highlighted in rectangles with dashed borders.

***H1***: *Task- and gaze-trained myoelectric interface performance improved significantly from early to late periods within trials*.

***H2***: *Task- and gaze-trained myoelectric interface performance in late period was not significantly different from mouse baseline*.

User performance in *Session 2* may be expected to differ from that of *Session 1* for two key reasons. First, users had more experience with the interface during the second session than the first, so although the decoders were initialized randomly the user may have adopted a different adaptation strategy. Second, users were explicitly instructed to fixate their gaze on the target in the second session. Therefore we tested the effect of session order on task performance achieved with the two training conditions with the last 20 seconds (*late*). To simultaneously assess the effect of training condition and session on performance, we used a two-way repeated measures ANOVA test with *α* = 0.05. We used paired t-tests with Bonferroni correction for post-hoc comparisons. We had the following two hypotheses for assessing the effect of interface conditions and session orders:

***H3***: *Performance in Session 2 was better than that in Session 1*.

***H4***: *Task- and gaze-trained myoelectric interfaces performance was not significantly different in Session 1 or Session 2*.

## 5 RESULTS

***H1***: *Task- and gaze-trained myoelectric interface performance improved significantly from early to late periods within trials*.

The tracking performance of both task-trained and gaze-trained interfaces improved throughout the three-minute training; in the *late* period, the users were able to control their cursors to approach the target as compared to *early* of a trial (Fig. 3). Paired t-tests found significant improvement in task performance from *early* to *late* periods for both interfaces (task-trained: *t* (10) = 4.93, *p <* 0.001; gaze-trained: *t* (10) = 3.40, *p <* 0.01). We visualized the 5th, 25th, 50th, 75th, and 95th percentiles of data distributions in boxplots (Fig. 4).

**Fig. 4.**
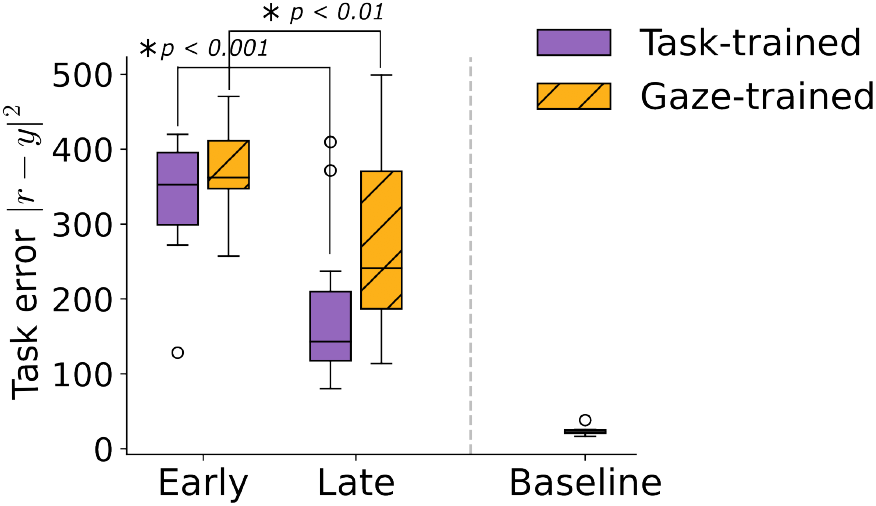
Boxplots (N = 11) of task performance in *early* and *late* periods, averaged across all sessions. The boxplot of the mouse baseline performance was shown for comparison. Lower task error corresponds to better task performance. Significant differences were labeled with * and *p* values (paired t-tests).

***H2***: *Task- and gaze-trained myoelectric interface performance in late period was not significantly different from mouse baseline*.

A repeated-measure ANOVA test showed significant differences across the three groups (*late* task-trained, *late* gaze-trained, and baseline) (Fig. 4; *F*_2,30_ = 20.64, *p <* 0.001). Post-hoc tests with Bonferonni correction indicated significant differences between *late* task-trained and baseline (*t* (10) = 8.34, *p <* 0.001), and between *late* gaze-trained and baseline (*t* (10) = 4.38, *p <* 0.01). However, it did not find a significant difference between *late* task-trained and *late* gaze-trained (*t* (10) = *−*2.51, *p* = 0.03), where the *p* value was above the corrected significance level.

***H3***: *Performance in Session 2 was better than that in Session 1*.

***H4***: *Task- and gaze-trained myoelectric interfaces performance was not significantly different in Session 1 or Session 2*.

The two-way repeated measure ANOVA test (sessions × interfaces) found a main effect for interface conditions but not session orders on task performance (Table 1). Although the ANOVA test did not find a significant effect on session orders, we observed that participants exhibited slight improvements from *Session 1* to *Session 2* (Fig. 5). Post-hoc tests with Bonferroni correction were then conducted to compare the interfaces in each session. Post-hoc tests indicated no significant difference between task- and gaze-trained interfaces in either session, with both *p* values above the corrected significance level (Fig. 6; *Session 1*: *t* (10) = *−*2.11, *p* = 0.061; *Session 2*: *t* (10) = *−*2.51, *p* = 0.031).

**Table 1.** Two-way repeated measure ANOVA (sessions × interfaces) results for time-domain error during testing.

| Factor | $F_{1,40}$ | $p$-value | Partial $\eta^2$ |
| --- | --- | --- | --- |
| Sessions | 1.40 | 0.24 | 0.03 |
| Interfaces | 5.17 | 0.028 | 0.11 |
| Interaction | < 0.001 | 0.99 | < 0.001 |

**Fig. 5.**
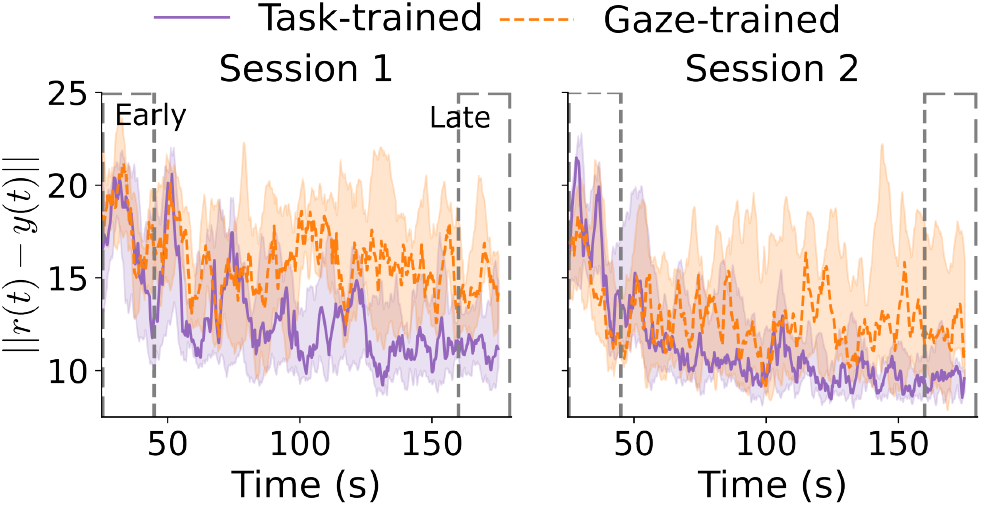
Distribution (median, interquartile, N = 11) of time-domain task error, | |*r − y* | |, of (a) *Session 1* and (b) *Session 2*, removing the initial 20-second ramp-up. Lower task error corresponds to better task performance. The *early* and *late* 20-second windows are highlighted in rectangles with dashed borders.

**Fig. 6.**
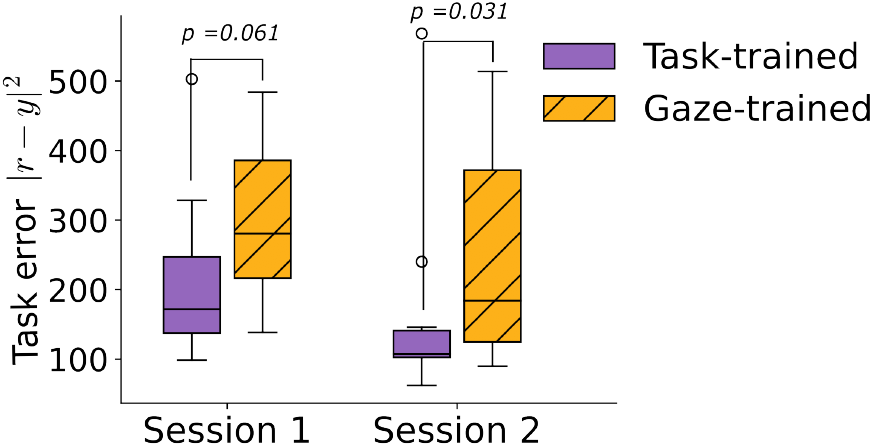
Boxplots (N = 11) of task performance in the *late* period, separated into *Session 1* and *Session 2*. Lower task error corresponds to better task performance. Comparisons between interfaces are shown with *p* values (paired t-tests).

## 6 DISCUSSION

This work used eye gaze to train an adaptive myoelectric interface for computer cursor control. The interface was adapted continuously based on real-time sEMG inputs and gaze data while users performed a continuous tracking task. We evaluated the performance of this adaptive myoelectric interface by comparing our novel gaze-trained to the previous task-dependent method [42] and investigated the effect of guided instruction on performance.

### 6.1 Task Performance Improved for Both Gaze-Trained and Task-Trained Interfaces

Our results demonstrate that both gaze- and task-trained interfaces worked effectively for training the adaptive myoelectric interface (Fig. 3; Fig. 4). This is consistent with the findings in Madduri et al. [42] and Aronson et al. [3], where the former showed that their task-trained adaptive interface improved tracking performance within minutes of training and the latter showed that natural gaze assisted in goal predicting for device control. However, as expected, the conventional mouse interface enables significantly better tracking than both myoelectric interfaces, underscoring the challenges in designing personalized myoelectric interfaces with high-dimensional inputs. Nevertheless, this study serves as a proof-of-concept, showcasing the feasibility of training an adaptive myoelectric interface using a task-agnostic approach.

### 6.2 Gaze-Trained Interface Performed Comparably to Task-Trained Interface

The two-way repeated measure ANOVA found a significant difference in interfaces (Table 1; *p* = 0.028); while the post-hoc tests further showed that the gaze-trained interface performed equally well with the task-trained interface (Fig. 6). The ANOVA result is not surprising as access to task information yields higher performance; however, the post-hoc tests demonstrate that the gaze-trained paradigm holds potential for real-world deployment even when the algorithm is not privileged with knowledge of the task. The novel gaze-trained interface presents a task-agnostic advantage over the previous task-trained interface, offering the potential to adapt while users engage in diverse task operations.

A preliminary experiment suggests that the gaze-trained interface is not limited to the tracking task presented in this paper. In the Video Figure, we demonstrate the potential of training this interface on a different class of reference signals (a sequence of stationary targets) and its potential to generalize to an unstructured handwriting task.

### 6.3 Differences in Performance with Additional Instruction and Practice

We further investigated the effect of session orders on task performance for each condition. In *Session 2*, participants received additional instruction on gaze fixation and had undergone more practice than in the first session. We observed that performances of both interfaces improved in *Session 2*, with a slightly lower error shown in Fig. 5. This improvement could be due to more consistent user behavior as they learn a strategy to control an interface, leading to proficient control [24, 52]. However, the ANOVA test did not find a significant improvement in performance given the additional instruction or practice in *Session 2*; rather, the performance of the two interfaces was more separated in this session (Fig. 6). This notable difference in performance between the two interfaces could be due to several factors, such as a shift in user strategy prompted by the instruction leading to improved task performance. Additionally, issues such as eye-tracking headset slippage [36] could have contributed to reduced accuracy in gaze data, as the eye-tracking headset was only calibrated at the beginning of the experiment; these errors could potentially accumulate over time, resulting in lower performance in later gaze-trained trials.

Finally, the gaze-trained interface effectively operated without guided instruction on where to look (*Session 1*). Although additional studies were still needed to know whether guided instruction affects performance, the gaze-trained interface surprisingly did not have a pronounced improvement when natural gaze was replaced with intentional gaze (*Session 2*). These findings highlight the robustness of our training paradigm and suggest that instruction on where to look might not be necessary for gaze-trained interface; instead, general practice across sessions or guidance in strategies might be sufficient to account for the improvement between sessions. Moreover, different from other gaze-controlled devices that use gaze as a direct control input [44, 53, 64], this approach allows natural eye movements during computer operations and can potentially minimize fatigue associated with intentional eye-tracking [34].

### 6.4 Potential Applications in HCI

#### This work is a proof-of-concept of a novel method

training an adaptive myoelectric interface with the user’s natural gaze. Imagine a wearable device that a user can inattentively put on their arm that automatically calibrates as they perform various tasks or switch between tasks in a virtual environment. Our work is a step toward realizing this vision, laying the groundwork for future investigations into continuously adaptive high-dimensional interfaces across a variety of HCI applications. We conclude by listing consumer and health applications that can potentially benefit from our interface.

#### Wearables in Augmented/Virtual Reality

Our gaze-trained myoelectric interface has the potential to enable cursor control when eye-tracking is available, such as using hand movements to navigate tabs in VR or swiping through slideshows in virtual presentations. For example, consider a user controlling a continuous cursor to navigate a web page in VR using a sEMG-based wearable device and a headset. Over time, the user may begin to sweat, feel fatigued and opt for smaller movements, or switch to another task, such as writing with the continuous cursor. Each of these scenarios leads to changes in sEMG signals. If using a conventional myoelectric interface, the user would need to perform another offline calibration to accommodate such signal changes. In contrast, our interface provides continuous adaptation on the fly, eliminating the need for offline re-calibration or prescribed tasks. This holds promise for seamlessly transitioning between user-led activities using user-selected hand movements.

#### Assistive Devices for Motor Control

While we tested our interface with healthy participants, the insights gained from this study have implications for assistive devices to restore motor abilities in healthcare settings. For example, our approach can potentially benefit sEMG- or electroencephalogram (EEG)-controlled wheelchair [2]. As gaze reveals a user’s intended moving direction, we can use gaze information to train an adaptive interface to control the continuous navigation of a wheelchair. This can be a promising alternative to current powered wheelchairs, which lack adaptability to individual users. Unlike previous gaze-controlled wheelchairs [45, 64], this approach integrates high-density biosignals to control the interface and does not require intentional gaze.

Another potential application lies in powered prostheses or robotic limbs controlled with high-density measurements like sEMG, EEG, or intracortical activities. Myoelectric prostheses still face high rates of abandonment, partly due to poor ease of use [18]. If eye information is accessible, our gaze-trained paradigm could be used to train an adaptive interface to control those devices, thus enhancing the usability of myoelectric prostheses across a diverse set of users. It is important to note that our study focused only on 2D cursor navigation. To extend its utility to the above applications, we will need to modify and validate the paradigm for 3D and multi-degree-of-freedom robotic limb control. Additionally, this will require depth information into eye-tracking data to estimate user intention in 3D.

#### 6.5 Limitations and Future Work

Our immediate next step is to expand the preliminary experiment showcased in the video and validate the applicability of our gaze-trained paradigm to less structured tasks. Specifically, we plan to train the adaptive interface while novice users engage in activities such as browsing web pages or writing with a cursor. Another future step is to incorporate the dynamic nature of eye movements in our gaze-trained paradigm. Eye movements have been shown to evolve as people learn novel sensory-motor mappings [12, 59] and are different between novice and skilled users [22]. This suggests that eye-tracking data will have additional benefits for adaptive interfaces beyond the training tested here. For instance, eye-tracking can be particularly helpful in detecting gradual user learning or abrupt shifts in user strategy, thereby facilitating prompt re-calibration of our myoelectric interface.

One limitation of this study was the limited number of subjects. Moreover, we only tested participants without disabilities. As myo-electric and eye-tracking devices hold promise for improving device accessibility for people with motor disabilities [69], our plan moving forward is to test the gaze-trained interface with individuals who have upper-limb motor disabilities, such as stroke survivors.

## 7 CONCLUSION

In this work, we propose a new method for training an adaptive myoelectric interface online using natural eye movements, treating the user’s gaze as a proxy for the task goal. Our gaze-trained approach eliminates the need for explicit task information in training myoelectric interfaces, offering a less supervised alternative compared to other methods. This approach also does not require users to be guided on where to focus their gaze, preventing drawbacks associated with intentional eye-tracking. This adaptive myoelectric interface holds the potential to extend to diverse tasks, including less structured computer operations with undefined task goals. This study will inform our ongoing work in developing an out-of-the-box myoelectric interface that adapts during diverse task operations, contributing to a more user-friendly and accessible class of interfaces.

## ACKNOWLEDGMENTS

We thank our participants in this study. This work was funded by Meta Reality Labs.

## A EYE DATA PRE-PROCESSING

To accurately estimate the task goals or the task targets from gaze, we pre-processed the raw gaze data using the following steps:

1. We found that raw gaze delayed the target reference by approximately 250-300 ms and had a phase lag of 0.4 radians below 1 Hz when a participant intentionally gazed at the target. This roughly aligned with the human visuomotor delay in a first-order tracking task [54]. To estimate the target position from gaze position, we removed the phase lag using the Fast Fourier Transform in the frequency domain.
2. After an initial calibration in PupilCapture (see section 4.2), we asked participants to fixate on nine static targets on the computer screen to examine any bias in gaze measurements. These biases were due to imperfect eye-tracker calibration or inaccurate transformation from camera coordinates to screen coordinates. We then removed biases in gaze positions in both horizontal and vertical axes.
3. We masked low-confidence eye measurements by removing gaze data below a confidence threshold of 0.5, in a range from 0 to 1 given by the PupilCapture Software. Low confidence gaze could be due to the user blinking or gazing off of the computer screen.

The cleaned gaze data had a significantly better correlation with the target trajectory *r* compared to the raw gaze data when a user was intentionally gazing at the target (Fig. 7).

**Fig. 7.**
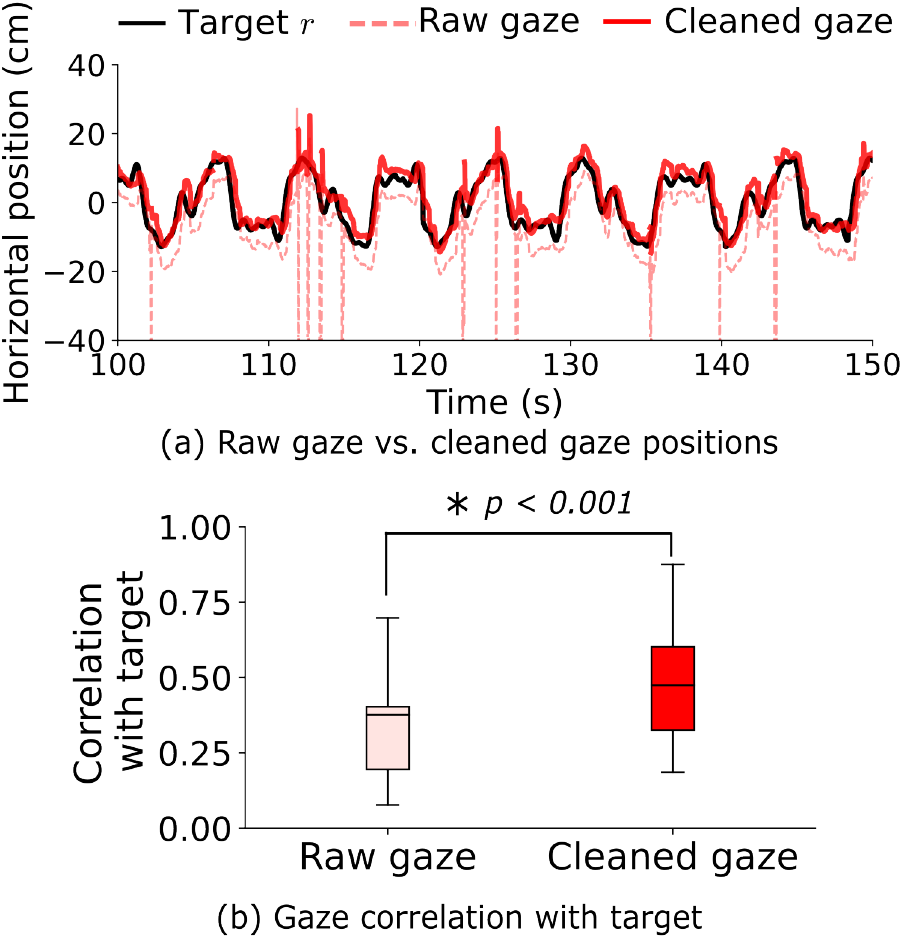
(a) An example of raw and cleaned gaze data when the user is gazing at the reference target *r*. Cleaned gaze data has fewer unwanted spikes compared to raw gaze data. (b) Boxplots (N = 11) of the Pearson correlation coefficient (range from 0 to 1) between raw gaze position and target *r* versus correlation coefficient between cleaned gaze position and target *r*, averaged across all trials. The significant difference was labeled with * and its *p* value (paired t-test with *α* = 0.05).

## B SUBJECTIVE USER EXPERIENCE

At the end of the experiment, participants provided subjective inputs, including the difficulty of the system (on a scale of 1*−*5, with 5 being the most difficult), the accuracy of the system (1*−*5, with 5 being the most accurate), and how much effort did it take to use the system (1*−*5, with 5 being a lot of effort). The median value of the three indices showed that participants rated the system to have relatively low (2) difficulty, slightly high (4) effort, and medium (3) accuracy (Fig. 8).

**Fig. 8.**
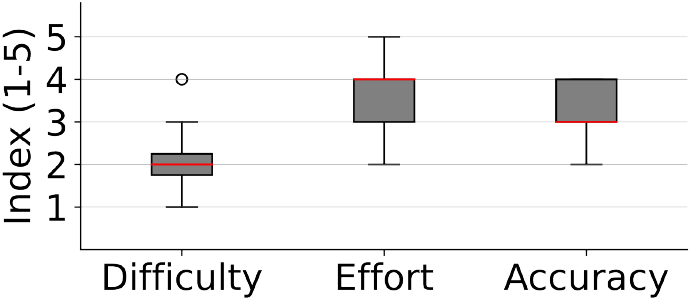
Boxplots (N = 11) of the difficulty, effort, and accuracy indices reported by the participants. The indices range from 1 to 5, with 5 being the most difficult, largest effort, and highest accuracy.

